# Opportunities to improve the recommendation of plant varieties under the Recommended List (RL) system

**DOI:** 10.1101/2024.06.07.597888

**Authors:** Chin Jian Yang, Joanne Russell, Ian Mackay, Wayne Powell

## Abstract

Recommended List (RL) is the UK plant variety recommendation system introduced in 1944 for supporting growers in making decisions on variety choices. The current analysis for RL is largely based on published works on trial designs and statistical analyses in the 1980s. Given the statistical advances that have been developed and adopted elsewhere, it is timely to review and update the methods for data analysis in RL. In addition, threats from climate change challenge the prediction of variety performance in future environments. Better variety recommendations, particularly for matching varieties to specific environments can be achieved through improved modeling of effects from genetics, environments and genetic by environment interactions. Here, we evaluated grain yield data from 153 spring barley varieties that were trialed for RL from 2002 to 2019. Our results show that the available methods for predicting variety performance are poor and inconsistent across environments without any clearly superior method. However, these shortcomings can be easily overcome by switching the statistical model for analyzing RL data to one that fits variety effects as random and accounts for genetic relationships among varieties. Lastly, we also discuss other possible approaches for analyzing RL data and highlight the relevance of genomics in both variety registration and recommendation.

## INTRODUCTION

Plant breeding, in combination with improved agronomy practices, contributes greatly toward advancing modern crop agriculture and safeguarding food security (Bradshaw 2017). Breeding programs are designed to produce new plant varieties with higher yield, qualities, disease resistance and abiotic stress tolerance (Cobb et al 2019). Much of the genetic gain in these traits can be attributed to either transgressive segregation in line breeding or heterosis with dispersed dominance in hybrid breeding (Mackay et al 2021). Because many target traits are polygenic and have complex genetic architecture, new varieties are produced through shuffling of favorable alleles or haplotypes within breeding pools (Scott et al 2021). For example, the Illinois Long-Term Selection Experiment for oil and protein content in maize grain has shown continuous genetic gains over more than a hundred years of selection (Moose et al 2004). Recent developments in precision technologies such as gene editing, remote sensing and artificial intelligence (Wallace et al 2018, Zaidi et al 2019, Xu et al 2022) could potentially revolutionize future breeding systems to achieve resilient, sustainable and regenerative agriculture.

As new varieties are released annually by various public and private breeding programs, plant variety recommendation system helps growers in choosing ideal varieties that fit their market goals and farm environments (Federizzi et al 2012). The variety recommendation system typically depends on analyses of trials that span multiple sites and years to characterize agronomic trait stability and rank across environments. In addition to new varieties, several important and high-performing older varieties are often included as checks or controls. Variety recommendation is usually limited to major crops with high economic importance due to the cost of large-scale trials. This system varies from country to country, which is known as Recommended List (RL) in the United Kingdom, National Variety Trials (NVT) in Australia, and Official Variety Testing (OVT) in the United States. RL is conducted by the Agriculture and Horticulture Development Board (AHDB) with the support from the Biomathematics and Statistics Scotland (BioSS), NVT is conducted by the Grain Research and Development Corporation (GRDC), and OVT is often conducted at the state-level by land-grant public universities. Each year, these testing authorities are responsible for analyzing trial data and publishing variety recommendations in various forms such as printed or online tables, variety selection tools or mobile applications.

Analyzing variety performance in multi-environment trials can be challenging due to the presence of Genetic by Environment Interaction (GEI) effects observed in most traits (Patterson and Silvey 1980). There are usually multiple replicates in each trial site and at least 10 to 20 trial sites per year depending on the geographical areas covered by the variety testing authorities. Appropriate modelling of genetic, environment and GEI effects are crucial to ensuring good predictions on variety performance. Variety trial design and analysis are century-old topics and trace back to early work such as Fisher (1935), Yates (1933, 1936, 1960) and Yates and Cochran (1938). The first edition of RL was published by the National Institute of Agricultural Botany (NIAB) in 1944 with only 15 winter wheat varieties (AHDB 2024). Interestingly, variety recommendation systems in many countries build on the approach developed by Patterson and Silvey (1980). UK was among the early adopters of this approach and has not changed its RL system since then. On the other hand, Australia has been constantly updating its NVT system to keep up with the state-of-the-art statistical developments (Smith et al 2001, 2005, 2015, 2021). In the US, the OVT systems vary from states to states and are rarely discussed in academic literature. In general, variety performance is usually obtained from a two-stage model where variety means are first computed for within environment (site-year) and followed by across environment. In some cases, a one-stage model is used where raw data from all trialed environments and replicates are fitted together. If appropriate variance components from the first stage are accounted in the second stage, the two-stage model is equivalent to the one-stage model (Piepho et al 2012).

Here, we evaluate the current RL system in the UK and identify possible areas for improvement. Performances of control and new varieties are often presented in a table of trait means within environment, where an environment is defined as site-year combination. This table is presumably derived from the approach of Patterson and Silvey (1980), but the information is not publicly available. Individual site, regional or overall means from one to five years of trials are calculated using simple averages regardless of missing environment. For our investigation, we choose spring barley as an example due to its importance in the UK agriculture and similarity to the RL system in other major crops such as winter barley, spring/winter wheat, spring/winter oats, and winter oilseed rape. We focus on grain yield because it has the most complete data in the public domain. We set the trial data in the immediate subsequent year as the prediction target. First, we evaluate the performance of current system across trial sites and years. Second, we test if the inclusion of genomic information and use of mixed models can help improve the system. Third, we estimate the annual yield loss based on the difference in performance between best predicted variety and best observed variety. Finally, we discuss opportunities for further improvement to take advantage of the latest developments in genetics, statistics and artificial intelligence.

Briefly, our results highlight key findings in several areas. First, the current RL system provides poor and inconsistent variety recommendations across environments. Second, while simple means from more trial sites lead to higher prediction accuracies on variety performance, one-year means outperform five-year means due to inaccurate variety ranks involving new and old varieties. Third, alternative methods using Genomic BLUP (GBLUP) address the shortcomings in the existing variety selection methods by producing better and more consistent variety performance predictions. Fourth, among eight different methods evaluated here, GBLUP with five-year of variety trial data outperforms all considered metrics. Fifth, the use of genomic marker data for variety recommendation fits perfectly with a previously proposed system for variety registration known as genomic Distinctness, Uniformity and Stability (DUS) (Yang et al 2021).

## MATERIALS AND METHODS

### Data collection

Treated yield data for spring barley was downloaded from the AHDB website (https://ahdb.org.uk/knowledge-library/recommended-lists-archive). The data included 153 unique varieties, 18 years (2002 to 2019) and 80 unique trial sites (Table S1, Table S2). The trial sites were further grouped based on “counties”, which were roughly defined based on 48 ceremonial counties in England, 32 council areas in Scotland, 22 principal areas in Wales and 6 counties in Northern Ireland (Figure S1, Table S1). Genomic marker data for all varieties were obtained from the IMPROMALT panel (Looseley et al 2020) which was genotyped using the barley 50k array (Bayer et al 2017). The original genomic marker data had 43,799 SNP markers and 809 varieties that span a much longer period of time. After filtering for the 153 varieties with yield data, we removed any monomorphic markers and retained 24,380 markers for analysis. Even though yield data was available for newer varieties from 2020 onward, their genomic marker data was unavailable and thus were not included in our analysis.

Historic data for 24 out of 37 weather stations were downloaded from the UK Met Office (https://www.metoffice.gov.uk/research/climate/maps-and-data/historic-station-data). These 24 weather stations were chosen based on their proximity to the trial sites. The data included monthly average of maximum temperature (C), minimum temperature (C), number of frost days and rainfall (mm) from 2002 to 2019. Temperature range (C) was estimated as the difference between the maximum and minimum temperatures.

The 2011 InFuse UK boundary data for local authorities was downloaded from the census website (https://borders.ukdataservice.ac.uk) for plotting the map. The local authorities in the boundary data were merged using the R/sf package (Pebesma and Bivand 2023) to form the counties. After merging, an approximated adjustment was applied to correct for part of county Armagh that was misclassified as county Down in Northern Ireland.

### Yield prediction

In the current UK variety selection tool, there are six possible methods for calculating variety performance to help growers with deciding on variety choice. These methods for making yield prediction in each target site were evaluated here and are henceforth referred to by their short names. I1: yield from the same trial site in previous one year. I5: yield mean from the same trial site in previous five years. R1: yield mean from all trial sites in the same region in previous one year. R5: yield mean from all trial sites in the same region in previous five years. A1: yield mean from all trial sites in previous one year. A5: yield mean from all trial sites in previous five years. Details of these methods are provided in Figure S2.

Two alternative methods for yield prediction, I5B and A5B, were evaluated by extending I5 and A5 with GBLUP approach, hence the names I5B and A5B. Instead of assuming that each variety was distinct, these alternative methods accounted for variety similarities by modelling trait variations according to their genomic relationships. I5B also accounted for random genetic by year interaction as GEI, but A5B did not. GBLUP is commonly used in breeding programs with genomic selection or prediction, and its predictive ability tends to scale with the number of individuals. In the case of variety trial, the number of varieties tend to be low and may not draw the full advantage of GBLUP. Due to the lack of replicated data within each trial site, we were unable to evaluate standard BLUP models which have been shown to be effective in improving prediction for variety recommendation (Piepho et al 2008). The models for I5B and A5B were fitted using the R/sommer package (Covarrubias-Pazaran 2018) based on the following general model equation.

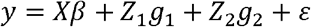

The model terms are described as below:

*y* is a vector of yield in *n* varieties.

*X* is a design matrix for the fixed effects such that the first column is a vector of 1’s for the overall mean, the second to fifth columns are vectors of 0’s and 1’s indicating which year the observation belongs to, the remaining columns are vectors of 0’s and 1’s indicating which trial site the observation belongs to.

*β* is a vector of fixed effects including the overall mean effect, year effects and site effects.

*Z*_1_ is an incidence matrix relating *n* varieties to observations *y*.

*g*_1_ is a vector of random genetic effect with a normal distribution of 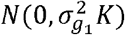.

*Z*_2_ is an incidence matrix relating *n* varieties by *p* years to observations y.

*g*_2_ is a vector of random genetic by year effect with a normal distribution of 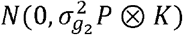.

*K* is an additive genetic relationship matrix in which the elements 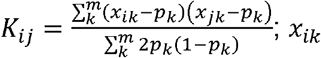 is the marker score for variety *i* at marker *k, X*_*jk*_ is the marker score for variety *j* at marker *k, p*_*k*_ is the allele frequency at marker *k*, and *m* is the total number of markers.

*P* is an identity matrix of *p* × *p*.

*ε* is a vector of residual effect with a normal distribution of 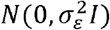 and is an identity matrix.

Specifically, the trial site effect was excluded in I5B and the genetic by year effect was excluded in A5B.

### Comparing predicted and observed yield

In practice, growers can choose a variety based on the performances of available varieties in previous years. Because we did not have the commercial production data for every variety in each site and year, we used the trial data as a proxy for prediction target. For example, yield prediction in 2007 was achieved using data from 2002 to 2006, and the predicted yield was compared to observed trial yield in 2007. Additionally, our criteria for analysis required a site to have trial data in the target year, the year immediately before, and a minimum of three out of five previous years (Figure S2).

Predicted yield from all eight methods was compared to the observed yield in each trial site and year using Spearman’s rank correlations. Given the importance of variety ranks in a variety recommendation system, ranked-based correlations were used instead of standard Pearson’s correlations. To ensure a fair comparison, the results were restricted to trial sites with available predicted yield from all eight methods. Because of our interest in the methods for identifying variety performance, the correlations were grouped according to method, method-by-year, or method-by-site. For each grouping approach, differences in means were tested using t-tests.

The consequences of sub-optimal variety choices due to imperfect predictions were quantified as Percent Deficit (PD) in yield for each method, site and year. To calculate PD, we first obtained the ratio of the difference between the highest and chosen variety’s yield over the difference between the highest and lowest yield. The ratio was then converted into percentage. For example, if the chosen variety’s, highest and lowest yields are 8, 10 and 5, respectively, then the PD would be 100*(10-8)/(10-5) = 40. All of the values used in the calculations were the observed yield in the target site and year.

### Data availability

R scripts, variety data (variety name, site, year, yield) and historical weather station data are available at https://github.com/cjyang-work/RL. Genomic marker data are available upon request from the authors in Looseley et al (2020). All online links have been archived at http://web.archive.org/ on 10 May 2024.

## RESULTS

### Trial environment

Between 2002 and 2019, 39 out of 108 counties in the UK had at least one spring barley variety trial for RL (Figure 1A, Figure S1, Table S1). Counties in the UK differ among the devolved nations and the definitions for counties used here are described in the Materials and Methods. The numbers of sites per county range from 1 to 5 with Perth and Kinross having the most sites (Aberuthven, Coupar Angus, Invergowrie, Kinross, Perth). Across the 39 represented counties over 18 years, the numbers of years with trials range from 1 (City of Edinburgh, Lancashire, Staffordshire) to 18 (Aberdeenshire, Fife, Norfolk). Similar to the wheat variety trial (Raymond et al 2023), trial sites for spring barley varieties may not always overlap from one year to another. Overall, spring barley variety trials can be found more often on the east coast than central or west coast of the UK (Figure 1A).

**Figure 1.**
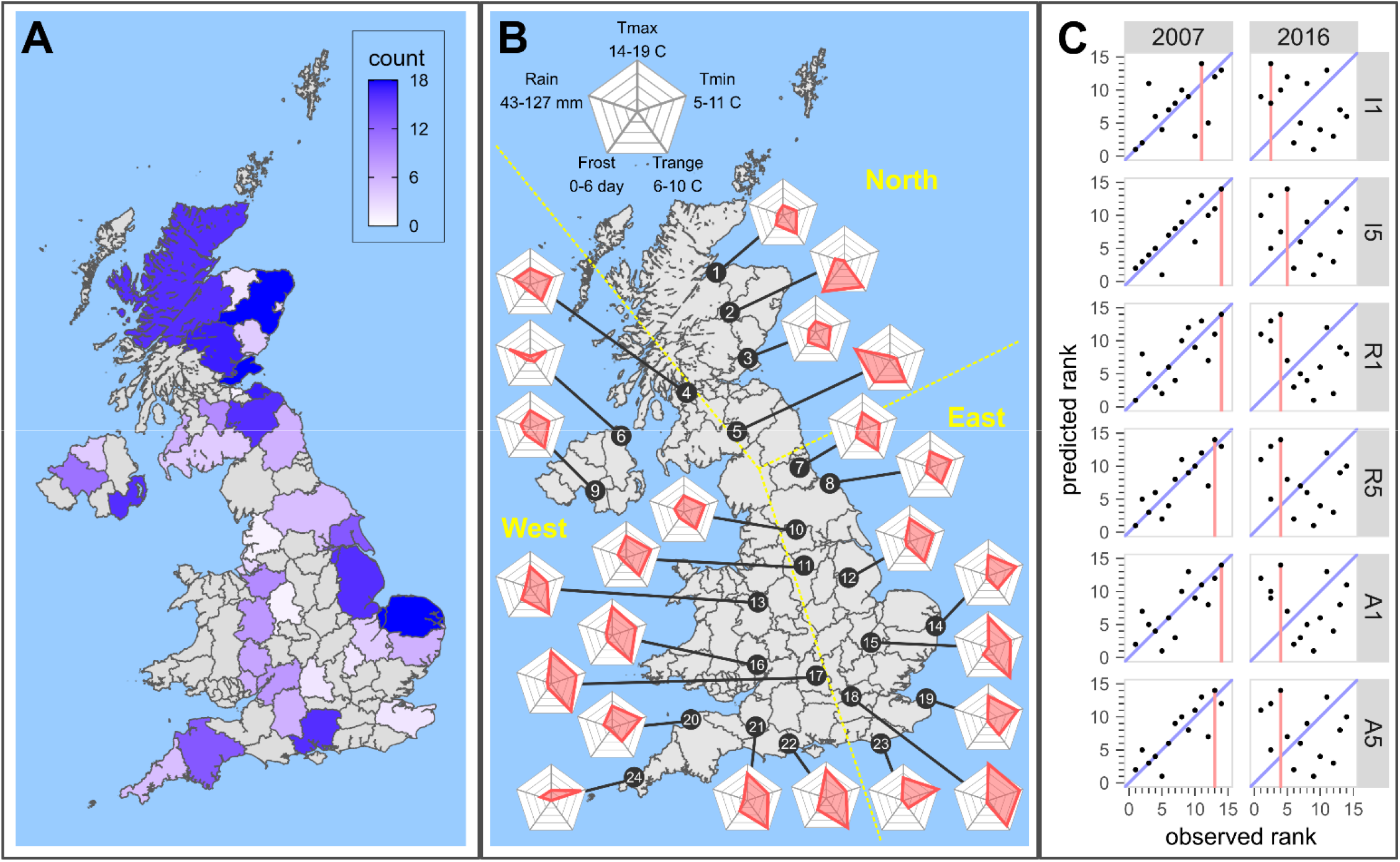
Trial distribution, climate variation and yield prediction. [**A**] Geographical distribution of UK counties with spring barley trials. [**B**] Climate variation from north (1) to south (24) and separation of trial sites into three regions (North, West, East). [**C**] Predicted versus observed yield ranks from six methods in 2007 and 2016 at Coaltown of Balgonie, which is located in Fife, Scotland and close to weather station 3.

The variety trials are spread across the key areas where spring barley is commonly produced, and these trial sites cover a broad range of climate variations. Historic data (2002 to 2019) from 24 relevant UK Met Office weather stations captures information on temperature, frost days and rainfall (Figure 1B, Table 1). Because the spring barley growing season typically ranges from April (or March if sown early) to August, we restrict the weather data from March to August. As expected, the maximum and minimum temperatures increase as latitude decreases. The switch from colder to warmer climate can be roughly placed between station 10 and 11, which is approximately close to a region known as the Midlands. Smaller temperature range is observed in coastal stations (example with lowest range) over inland stations (example with highest range). The risk of frost during spring barley season is fairly low, but there are several north and inland stations with higher numbers of frost days such as 2 and 5. Rainfall is evenly distributed across the stations except for three stations in the northwest UK (4, 5, 6) with higher rainfall.

**Table 1.** Weather station data. Out of 37 weather stations with archived records from the UK Met Office, 24 with close proximity to spring barley trial sites are shown here. For each weather station, temperature (maximum/minimum/range), number of frost days and rainfall are provided as means from March to August of 2002 to 2019.

| ID | Weather Station | Longitude (°) | Latitude (°) | T<sub>max</sub> (°) | T<sub>min</sub> (°) | T<sub>range</sub> (°) | Frost (day) | Rain (mm) |
| --- | --- | --- | --- | --- | --- | --- | --- | --- |
| 1 | Nairn | -3.821 | 57.593 | 15.26 | 7.05 | 8.20 | 1.93 | 56.80 |
| 2 | Braemar | -3.396 | 57.011 | 13.95 | 4.88 | 9.07 | 5.48 | 66.70 |
| 3 | Leuchars | -2.861 | 56.377 | 15.36 | 7.30 | 8.06 | 2.02 | 61.44 |
| 4 | Paisley | -4.430 | 55.846 | 16.13 | 8.33 | 7.79 | 1.09 | 83.03 |
| 5 | Eskdalemuir | -3.205 | 55.312 | 14.41 | 5.93 | 8.48 | 3.93 | 129.14 |
| 6 | Ballypatrick Forest | -6.153 | 55.181 | 13.84 | 7.56 | 6.28 | 0.88 | 96.95 |
| 7 | Durham | -1.585 | 54.768 | 16.19 | 7.68 | 8.51 | 1.63 | 56.65 |
| 8 | Whitby | -0.624 | 54.481 | 16.10 | 8.32 | 7.77 | 1.04 | 52.10 |
| 9 | Armagh | -6.649 | 54.352 | 16.34 | 7.97 | 8.37 | 1.43 | 67.24 |
| 10 | Bradford | -1.772 | 53.813 | 16.19 | 8.32 | 7.87 | 1.23 | 66.22 |
| 11 | Sheffield | -1.490 | 53.381 | 16.97 | 8.93 | 8.04 | 0.81 | 65.04 |
| 12 | Waddington | -0.522 | 53.175 | 17.12 | 8.71 | 8.41 | 1.00 | 55.88 |
| 13 | Shawbury | -2.663 | 52.794 | 16.91 | 7.66 | 9.25 | 2.20 | 54.92 |
| 14 | Lowestoft | 1.727 | 52.483 | 16.81 | 9.48 | 7.32 | 0.71 | 49.73 |
| 15 | Cambridge NIAB | 0.102 | 52.245 | 18.33 | 8.52 | 9.81 | 1.49 | 44.68 |
| 16 | Ross-On-Wye | -2.584 | 51.911 | 17.97 | 8.76 | 9.20 | 1.26 | 58.65 |
| 17 | Oxford | -1.262 | 51.761 | 18.51 | 9.05 | 9.46 | 0.97 | 53.87 |
| 18 | Heathrow | -0.449 | 51.479 | 19.16 | 10.04 | 9.11 | 0.64 | 46.79 |
| 19 | Manston | 1.337 | 51.346 | 17.35 | 9.68 | 7.67 | 0.48 | 46.52 |
| 20 | Chivenor | -4.147 | 51.089 | 16.96 | 9.62 | 7.35 | 0.66 | 65.27 |
| 21 | Yeovilton | -2.641 | 51.006 | 17.73 | 8.29 | 9.44 | 2.03 | 52.41 |
| 22 | Hurn | -1.835 | 50.779 | 17.98 | 8.21 | 9.77 | 2.25 | 57.35 |
| 23 | Eastbourne | 0.285 | 50.759 | 17.17 | 10.59 | 6.58 | 0.28 | 48.14 |
| 24 | Camborne | -5.327 | 50.218 | 15.28 | 9.83 | 5.45 | 0.22 | 69.87 |

### Variety selection tool

In the variety selection tool provided by the AHDB (https://ahdb.org.uk/variety-selection-spring-barley), users can select one or multiple traits and calculate the performance of all available varieties using six different methods. These methods generate variety performance based on simple means of the chosen trait(s) and the details are provided in the Material and Methods section. Briefly, the trait means can be calculated for each trial site (I), region (R) or all sites (A) from 1 or 5 years of data. For a fair comparison of variety selection methods, only trial sites with data in the target year, the year before, and a minimum of three out of five previous years are considered (Figure S2). These criteria results in 29 out of 80 trial sites analyzed (Table 2). AHDB classifies the regions for cereal into North, West and East, but these regions do not actually reflect the observed climate variations (Figure 1B, Table S1). For any chosen method, users are given ranks of the spring barley varieties from the best to worst. For simplicity, we are focusing on grain yield in our analysis because that is the most important agronomic trait and has the most publicly available data.

**Table 2.**
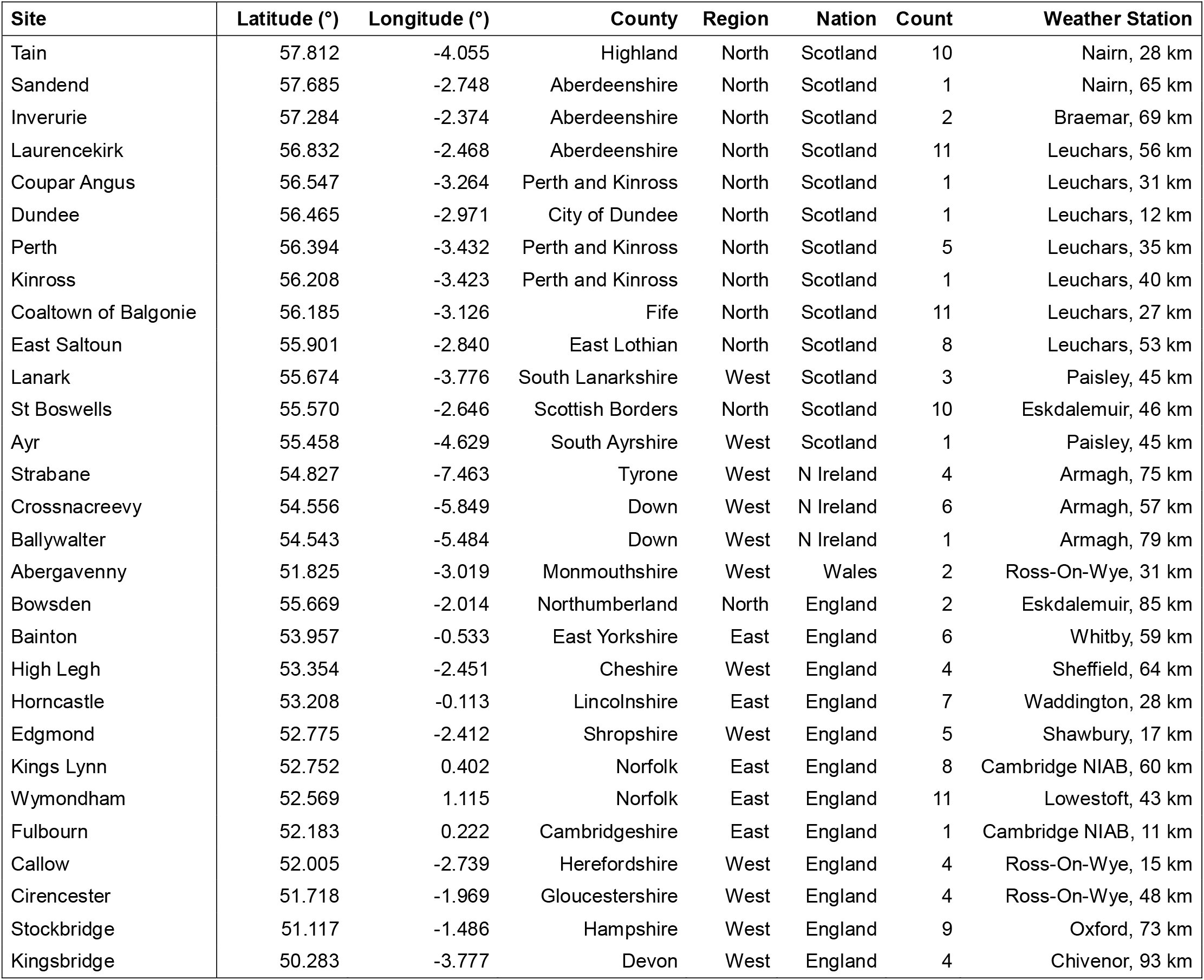
Summary of spring barley trial sites. Out of 80 unique trial sites, 29 with sufficient data have been included in our analyses. For each of these sites, additional geographical information, count of analyzed years and closest weather stations are provided.

| Site | Latitude (°) | Longitude (°) | County | Region | Nation | Count | Weather Station |
| --- | --- | --- | --- | --- | --- | --- | --- |
| Tain | 57.812 | -4.055 | Highland | North | Scotland | 10 | Nairn, 28 km |
| Sandend | 57.685 | -2.748 | Aberdeenshire | North | Scotland | 1 | Nairn, 65 km |
| Inverurie | 57.284 | -2.374 | Aberdeenshire | North | Scotland | 2 | Braemar, 69 km |
| Laurencekirk | 56.832 | -2.468 | Aberdeenshire | North | Scotland | 11 | Leuchars, 56 km |
| Coupar Angus | 56.547 | -3.264 | Perth and Kinross | North | Scotland | 1 | Leuchars, 31 km |
| Dundee | 56.465 | -2.971 | City of Dundee | North | Scotland | 1 | Leuchars, 12 km |
| Perth | 56.394 | -3.432 | Perth and Kinross | North | Scotland | 5 | Leuchars, 35 km |
| Kinross | 56.208 | -3.423 | Perth and Kinross | North | Scotland | 1 | Leuchars, 40 km |
| Coaltown of Balgonie | 56.185 | -3.126 | Fife | North | Scotland | 11 | Leuchars, 27 km |
| East Saltoun | 55.901 | -2.840 | East Lothian | North | Scotland | 8 | Leuchars, 53 km |
| Lanark | 55.674 | -3.776 | South Lanarkshire | West | Scotland | 3 | Paisley, 45 km |
| St Boswells | 55.570 | -2.646 | Scottish Borders | North | Scotland | 10 | Eskdalemuir, 46 km |
| Ayr | 55.458 | -4.629 | South Ayrshire | West | Scotland | 1 | Paisley, 45 km |
| Strabane | 54.827 | -7.463 | Tyrone | West | N Ireland | 4 | Armagh, 75 km |
| Crossnacreevy | 54.556 | -5.849 | Down | West | N Ireland | 6 | Armagh, 57 km |
| Ballywalter | 54.543 | -5.484 | Down | West | N Ireland | 1 | Armagh, 79 km |
| Abergavenny | 51.825 | -3.019 | Monmouthshire | West | Wales | 2 | Ross-On-Wye, 31 km |
| Bowsden | 55.669 | -2.014 | Northumberland | North | England | 2 | Eskdalemuir, 85 km |
| Bainton | 53.957 | -0.533 | East Yorkshire | East | England | 6 | Whitby, 59 km |
| High Legh | 53.354 | -2.451 | Cheshire | West | England | 4 | Sheffield, 64 km |
| Horncastle | 53.208 | -0.113 | Lincolnshire | East | England | 7 | Waddington, 28 km |
| Edgmond | 52.775 | -2.412 | Shropshire | West | England | 5 | Shawbury, 17 km |
| Kings Lynn | 52.752 | 0.402 | Norfolk | East | England | 8 | Cambridge NIAB, 60 km |
| Wymondham | 52.569 | 1.115 | Norfolk | East | England | 11 | Lowestoft, 43 km |
| Fulbourn | 52.183 | 0.222 | Cambridgeshire | East | England | 1 | Cambridge NIAB, 11 km |
| Callow | 52.005 | -2.739 | Herefordshire | West | England | 4 | Ross-On-Wye, 15 km |
| Cirencester | 51.718 | -1.969 | Gloucestershire | West | England | 4 | Ross-On-Wye, 48 km |
| Stockbridge | 51.117 | -1.486 | Hampshire | West | England | 9 | Oxford, 73 km |
| Kingsbridge | 50.283 | -3.777 | Devon | West | England | 4 | Chivenor, 93 km |

Predictions of variety performance are highly variable and vary in success across the years (Figure 1C). Using Coaltown of Balgonie, one of the most trialed sites, as an example, the predicted variety ranks are well correlated with the observed ranks in 2007 but not 2016. In 2007, the best prediction is achieved in method I5, and the best observed variety matches with the best predicted variety in methods I5, R1 and A1. When considering all varieties, five-year means generally perform better than one-year means. However, in 2016, none of the methods has a clear advantage over another as they all perform badly. The best predicted varieties end up with poor yield across all methods.

### Comparison across methods in predicting variety yield

The best prediction of variety performance across all trial sites and years can be achieved using A5B with a median correlation between predicted and observed variety ranks of 0.571 (Figure 2). The statistical model for A5B treats variety effect as random, accounts for the genetic relationships among varieties, and calculates the variety effect as GBLUP. The improvement in prediction with this approach is due to shrinkage toward means and maximization of true and predicted genotypic values (Piepho et al 2008). In addition, the A5B method takes advantage of the tendency of similar varieties to produce similar phenotypes. The next best prediction comes from both I5B and A1 with a median correlation of 0.518. While I5B uses a similar model to A5B, it uses only data from single trial sites. This discrepancy highlights the benefits of variety performance prediction from multiple over single trial sites. Curiously, I5 has the worst prediction with a median correlation of 0.324. Despite different methods showing different predictive abilities, the use of GBLUP offers a clear advantage over other methods.

**Figure 2.**
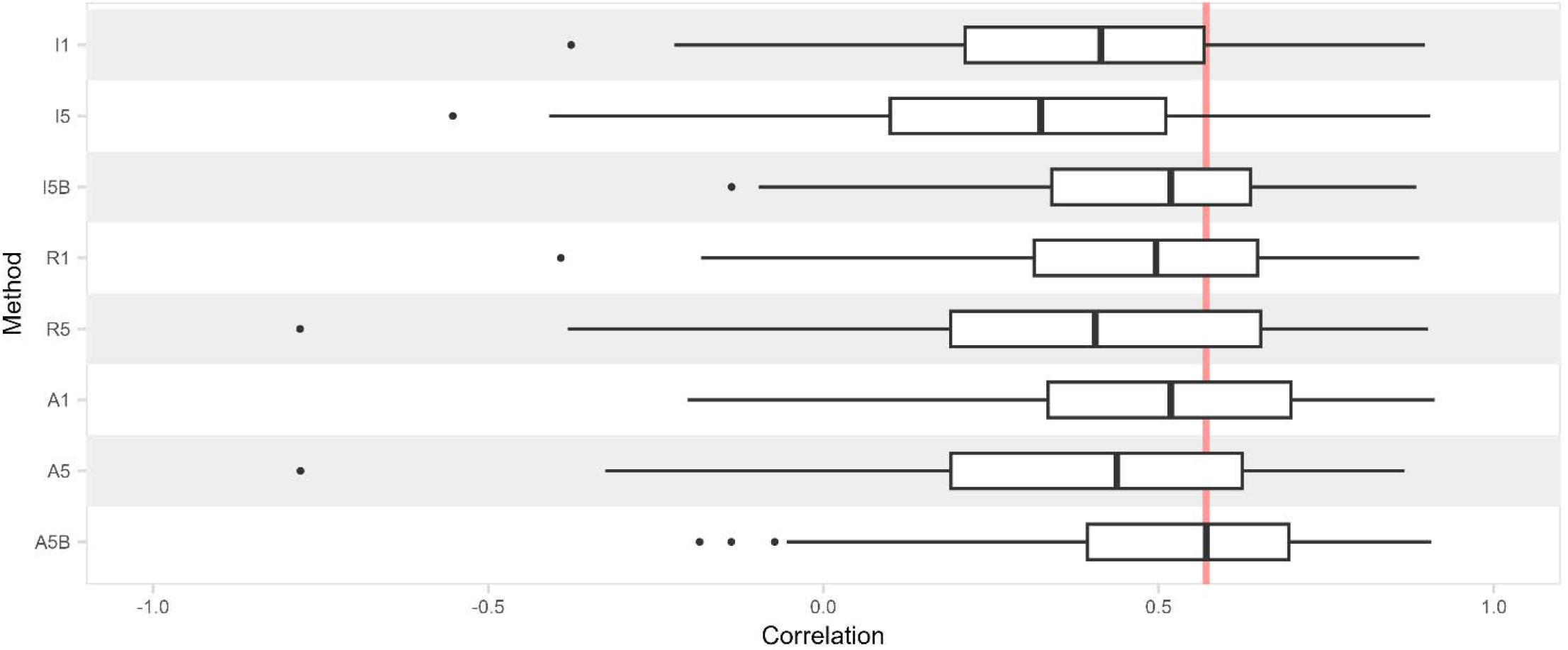
Comparison of eight different methods for predicting variety performance. Each boxplot contains correlations between predicted and observed yield ranks from 80 unique trial sites and 13 years (2007 to 2019). The lower, middle and upper hinges of the boxplots represent the first, second (median) and third quartiles of the data points. The whiskers extend from the boxes to the furthest data points within 1.5 times of the interquartile range. The vertical red line serves as a reference based on the largest median correlation from method A5B.

Methods that use data from more trial sites show higher accuracies in the prediction of variety performance. This pattern follows the order of yield means from individual sites (I), regional sites (R) and all sites (A) (Figure 2). The numbers of trial sites range from 2 to 11 per year for R and 13 to 25 per year for A (Table S1). The median correlations between predicted and observed variety ranks for I1/I5, R1/R5 and A1/A5 are 0.377, 0.474 and 0.488, respectively. The correlations for R-based methods are closer to the A-based methods instead of I-based methods, which could be due to diminishing benefits from additional trial sites and limitations of methods that derive variety performance through simple means.

In contrast to the GBLUP methods, the other methods work better with one year of yield data instead of five years. This observation is consistent across all three comparisons for the pairs of I1-I5, R1-R5 and A1-A5 (Figure 2). The general expectation is that better prediction of variety performance can be attained using more years just as shown by using more trial sites. There are two possible explanations for this contradiction. First, year-to-year yield variation can be large and data from years further apart are less informative toward predicting the yield in target year. Second, given the nature of variety recommendation system where new and old varieties constantly enter and exit each year, some varieties have fewer data than others in the predictions with five years of data and lead to an unbalanced estimate of variety means. These issues are partially overcome through the use of GBLUP methods, and likely explain the poor prediction of variety performance through simple means.

### Site-specific comparison of methods

Results for the six most used trial sites suggest a slight advantage for GBLUP methods (Figure 3). Four of these sites (Coaltown of Balgonie, Laurencekirk, St Boswells and Tain) are located in Scotland and two (Stockbridge, Wymondham) are located in England. Full results for all 29 analyzed trial sites are provided in Figure S3. The only comparison that is significant after applying Bonferroni correction for multiple testing is between method I1 and A5B (p = 0.0011), where A5B produces significantly higher correlations between predicted and observed variety ranks (Figure 3B). Despite the lack of significance in other pairwise comparisons, the GBLUP methods tend to perform better than their counterparts (e.g. I5B vs I5 and A5B vs A5) in all six trial sites.

**Figure 3.**
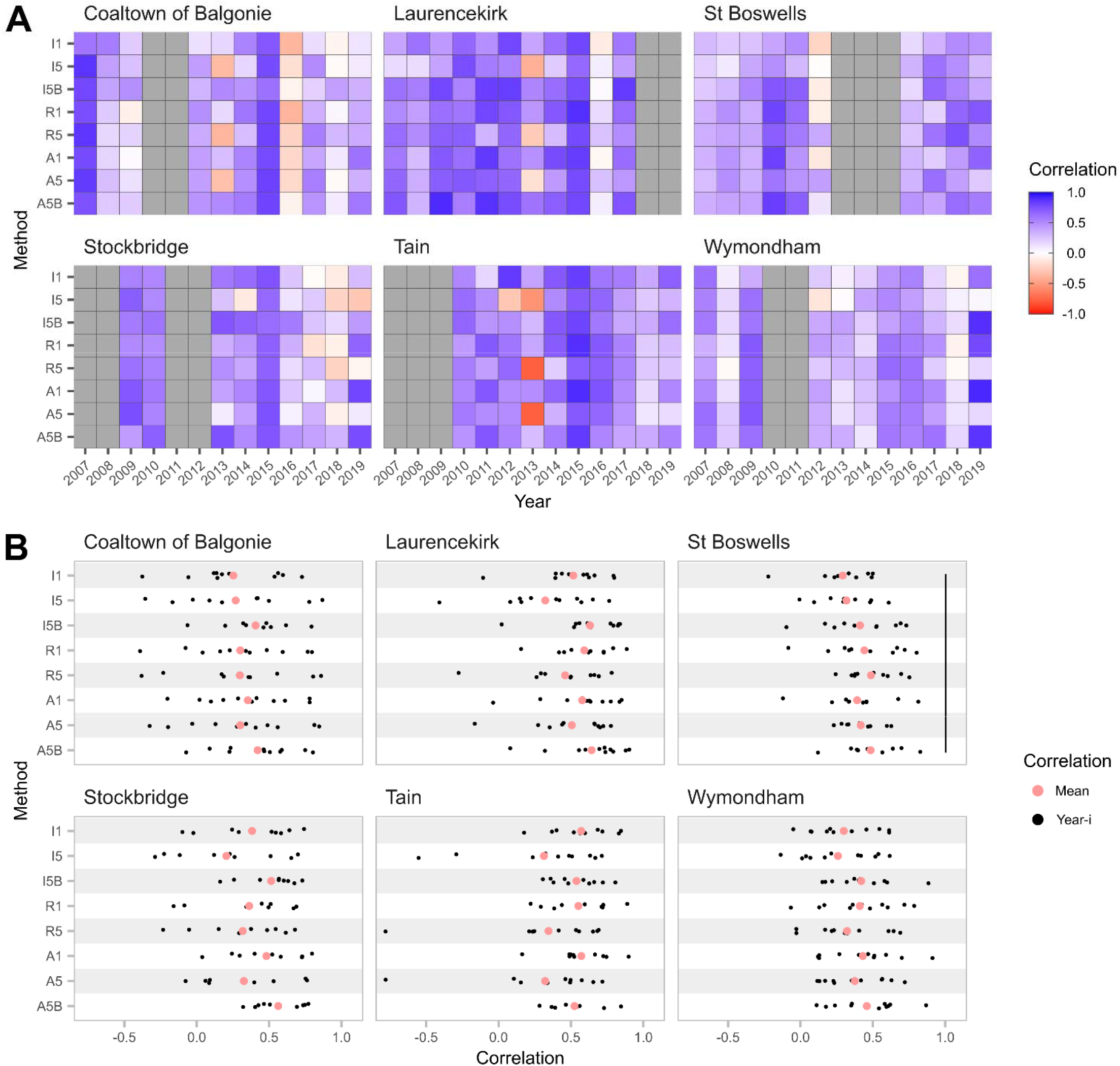
Correlations between predicted and observed yield ranks in six trial sites. [**A**] Heatmap of correlations for eight methods and 13 years. These six trial sites are highlighted here because they have the greatest number of trials across the years. [**B**] Scatter plot of correlations for eight methods and 13 years. Significant difference in mean correlations between methods after applying Bonferroni correction is indicated by a black vertical line.

A previous observation that non-GBLUP methods work better with one year of yield data instead of five years (Figure 2) can again be seen in Laurencekirk, Stockbridge and Tain (Figure 3). As an extreme example, method I5, R5 and A5 produce negative correlations when used to predict 2013 yield at Laurencekirk and Tain with data from 2008 to 2012 (Figure 3A). Similar examples can also be seen at Coaltown of Balgonie in 2013, Stockbridge in 2018 and 2019, and Wymondham in 2019. In many cases, GBLUP methods are able to recover the correlations to similar levels achieved from using one year of data. In contrast to several occasions where non-GBLUP methods produce negative correlations, GBLUP methods, especially A5B, offer better consistencies and lower risk of negative correlations.

Methods aside, varying success in yield prediction can be found across trial sites. Among the six trial sites, Laurencekirk appears to be the most consistent and best predicted over the years (Figure 3A) with a mean correlation of 0.708 across all methods and years. Tain places second with a mean correlation of 0.623. The remaining trial sites have mean correlations of 0.541 at St Boswells, 0.526 at Stockbridge, 0.496 at Wymondham and 0.434 at Coaltown of Balgonie. Correlations at Coaltown of Balgonie and Stockbridge drop in recent years while correlations at St Boswells and Wymondham are consistently low in most years.

### Year-specific comparison of methods

Good yield prediction can be achieved using GBLUP methods in all analyzed years from 2007 to 2019 (Figure 4). A5B slightly outperforms I5B but they are highly comparable. Each year, A5B and I5B either perform better than non-GBLUP methods or perform similarly to the best non-GBLUP method. Correlations for yield prediction with I5B are significantly higher than I5 (p = 0.0004) and A5 (p = 0.0002) in 2013, as well as I5 (p = 0.0002) and A5 (p = 0.0001) in 2019. Correlations for yield prediction with A5B are significantly higher than A5 (p = 0.0007) in 2013, A5 (p = 0.0007) in 2018, as well as I5 (p = 0.0003), R5 (p = 0.0008) and A5 (p < 0.0001) in 2019. These comparisons are significant after applying Bonferroni correction for multiple testing (Figure 4A). As found in the previous analysis on six most trialed sites, GBLUP methods are less likely to produce negative correlations on yield prediction.

**Figure 4.**
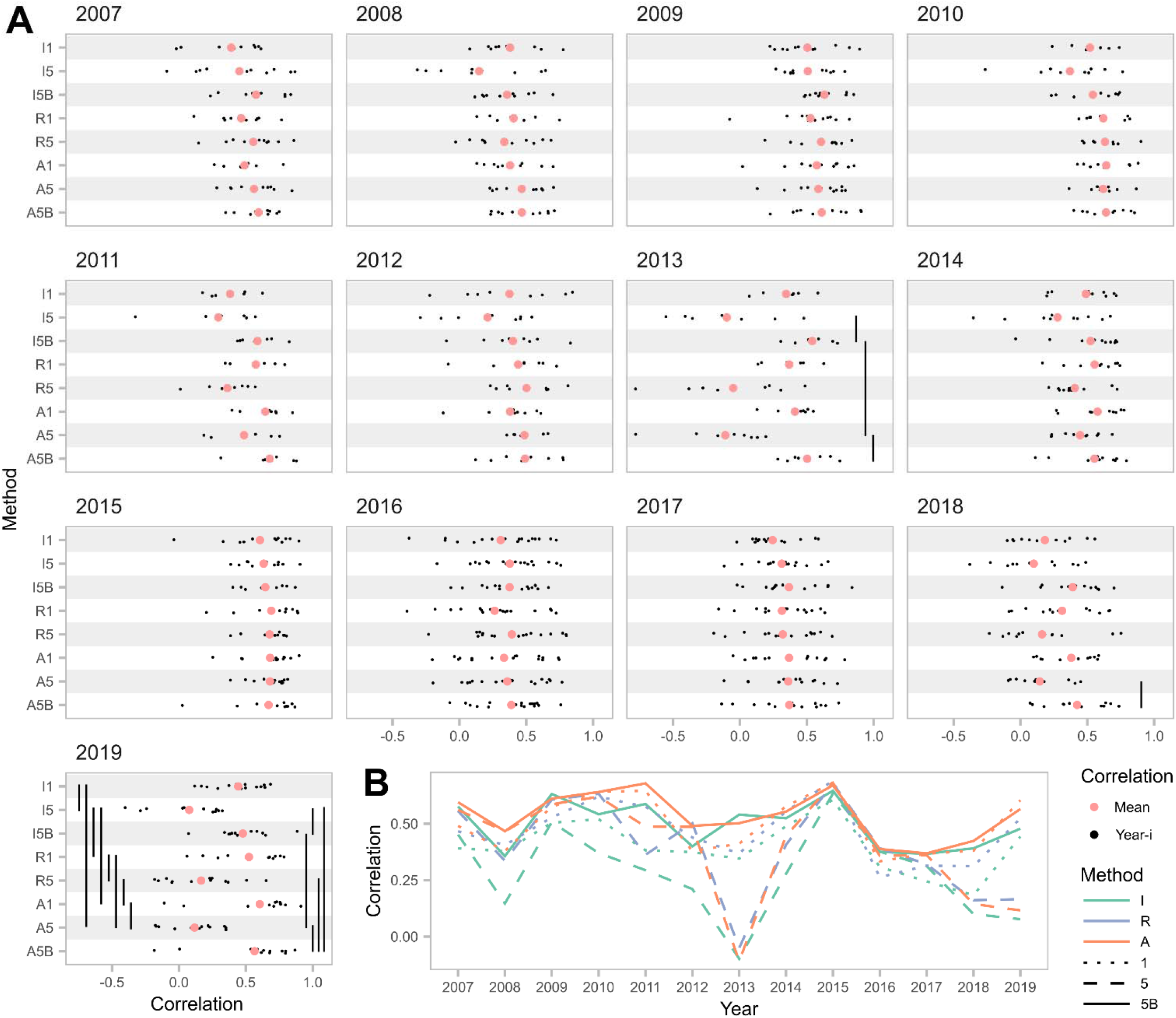
Correlations between predicted and observed yield ranks over 13 years. [**A**] Scatter plot of correlations for eight methods and trial sites within each year. Significant differences in mean correlations between methods after applying Bonferroni correction are indicated by black vertical lines. [**B**] Line plot of mean correlations for eight methods shown from 2007 to 2019.

In addition, GBLUP methods offer higher consistency than other methods over the years (Figure 4B). Correlations from I5B and A5B constantly hover close to 0.5 with smaller spikes and drop-offs. Non-GBLUP methods, especially I5, R5 and A5, suffer from severe correlation drops in few selected years. This observation again highlights the challenges of using five years of variety trial data due to bias against new varieties with less trial data and failure of existing methods in the variety selection tool to account for this bias. Among the three non-GBLUP methods that use five years of trial data, I5 clearly underperforms in most years. This result highlights the strong environment and GEI effects on yield in different years.

Methods aside, varying success in yield prediction can again be found across the years. The best year is 2015 with a mean correlation of 0.661 across all methods and trial sites. This is also the year where all methods produce similar correlations (Figure 4). Other good years with mean correlations above 0.500 are 2007, 2009, 2010 and 2011. In contrast, 2013 is the worst year for yield prediction with a mean correlation of 0.239. Poor yield prediction with mean correlations below 0.400 can be seen in 2008, 2016, 2017, 2018 and 2019. In most of the years where the yield prediction is poor, much higher correlations can be attained using GBLUP methods.

### Quantifying deficits in yield

Because of the imperfect yield prediction, all methods result in some amount of deficit in yield across trial sites and years (Figure 5, Figure 6). Percent Deficits (PD) in yield are calculated as the percentage of difference between best observed yield and chosen variety’s yield over the difference between best and worst observed yields. The chosen variety is taken from the best predicted variety using the eight methods. PD at trial sites can be hard to visualize on a map and are therefore merged into counties by averaging PD from all sites within the same county each year (Figure 5). Most counties exhibit moderate PD with small variation among the methods. Extreme PD in a few counties such as Northumberland (Bowsden site), South Ayrshire (Ayr site) and Cambridgeshire (Fulbourn site) needs to be treated with caution as they only have one to two years of predicted data for analysis. Overall, we observe a mean PD of 29 across all methods, trial sites and years.

**Figure 5.**
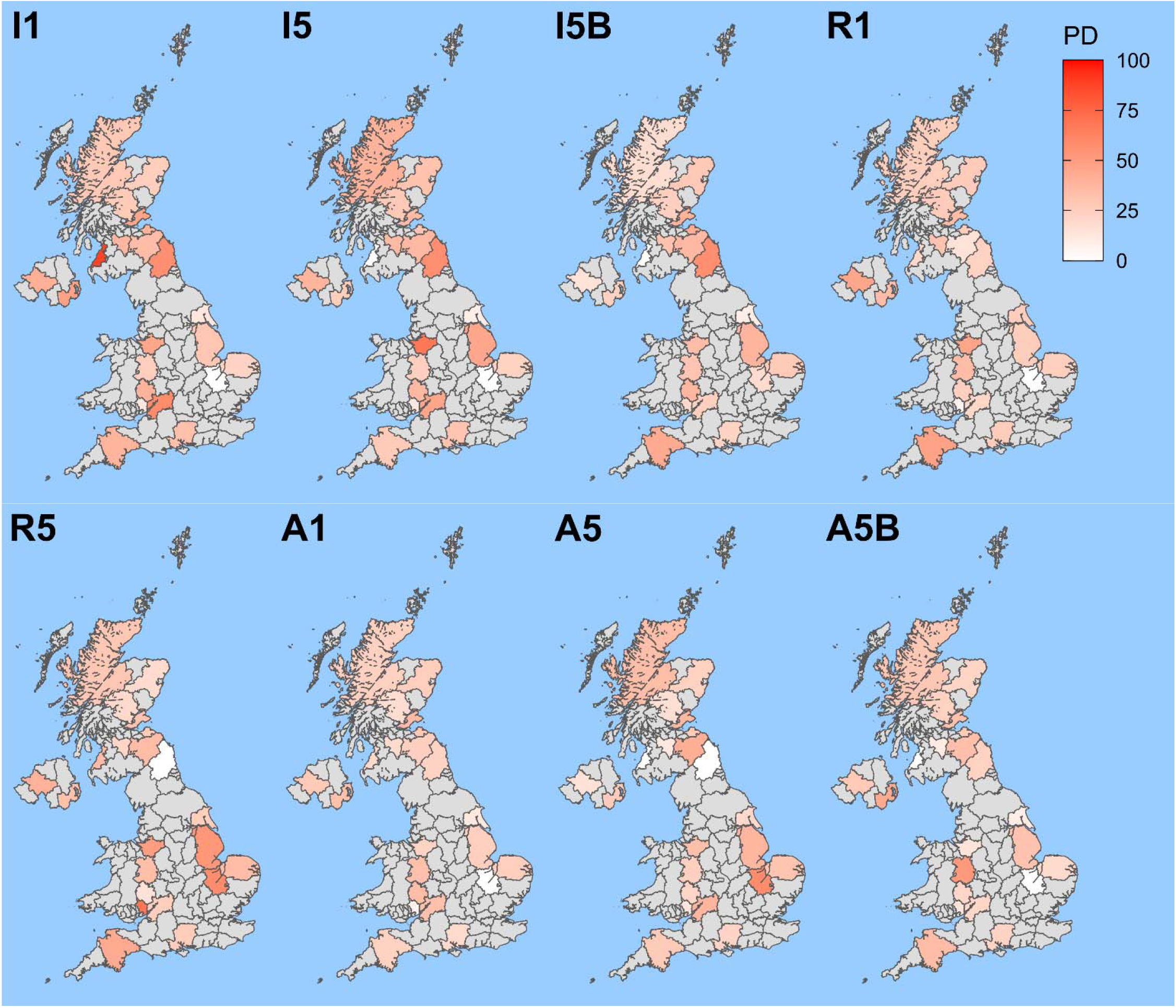
Distribution of yield deficits across counties. Percent Deficits (PD) in yield due to incorrect predictions of best local variety are quantified for each trial site and year. The heat map shows the county-level PD as the means across all trial sites and years within each county.

**Figure 6.**
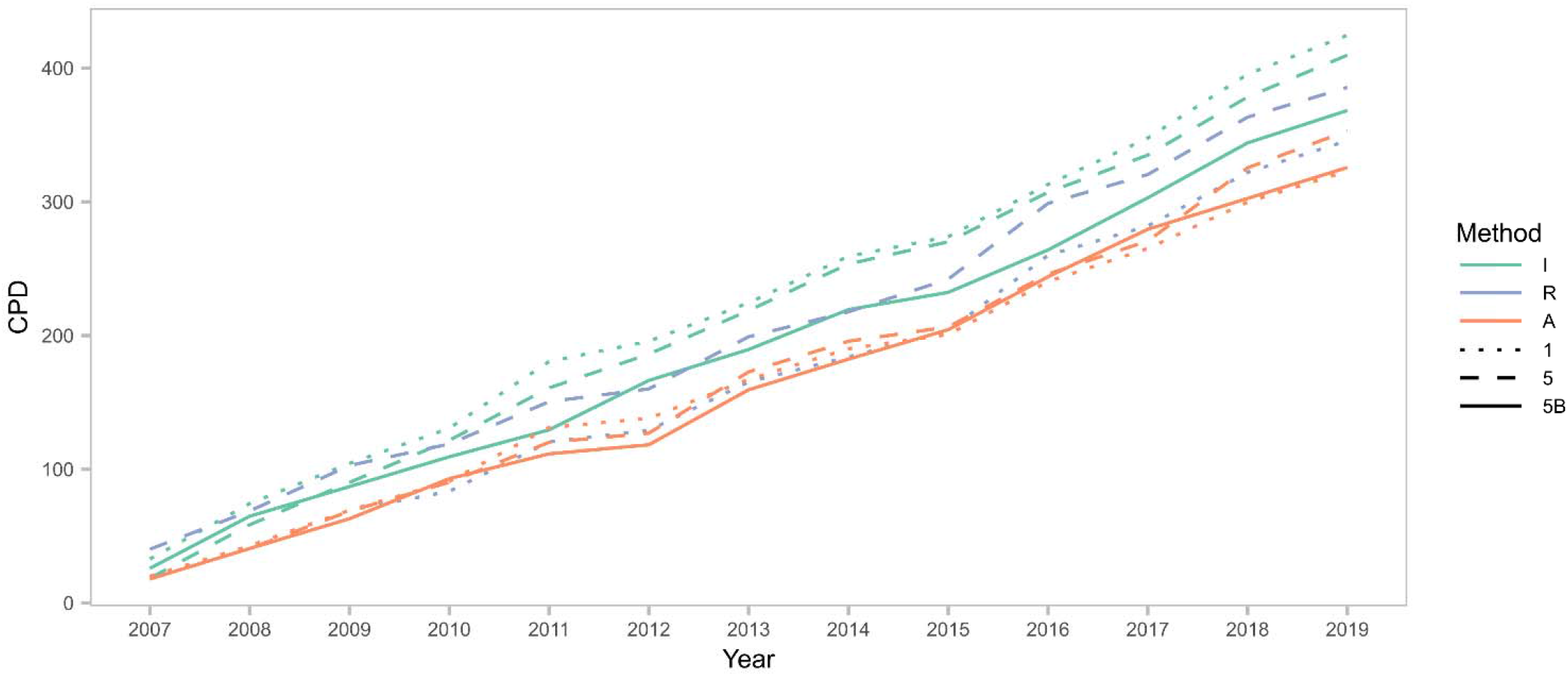
Cumulative Percent Deficits (CPD) in yield over the years. PD is averaged across all trial sites within each year to obtain the annual mean PD, which is then summed over the years from 2007 to 2019 to highlight the CPD in yield for each method.

While it can be hard to discern the PD from different methods at geographical level, we find consistent differences among the methods across the years. A1 and A5B produce the lowest Cumulative PD (CPD), followed by comparable CPD in A5 and R1 (Figure 6). On the other hand, high CPD in I1, I5 and R5 suggests that these methods are not suitable at predicting variety performance. Despite I5B and A5B having comparable correlations, I5B is mediocre in terms of CPD as it is largely hovering between the good and poor methods. Interestingly, there is a discrepancy of about 100 CPD between the best and worst method over 13 years. As a caveat, we have to assume that all sites contribute equally to the UK annual spring barley production in our estimation of CPD. While this assumption is unlikely to be close to the reality for any given year, it should still produce a reasonable estimate over a period of time as the trial sites are designed to cover spring barley production areas in the UK.

## DISCUSSION

The UK variety recommendation system builds on the work of Patterson and Silvey (1980) and has remained largely unchanged since then. While the trial design and analysis for RL are highly comprehensive at the level of individual trial site (https://ahdb.org.uk/ahdb-recommended-lists-for-cereals-and-oilseeds-2021-2026), the variety performance has not been well evaluated from a multi-environment perspective. The current method for analyzing variety performance largely focuses on estimating means at each trial site without accounting for phenotypic variations due to genetic relatedness among varieties, environmental effects and GEI across trial sites and years. Many examples highlight the importance of GEI in phenotypic variations (Williams et al 2008, Zhao and Xu 2012, Lopez-Cruz et al 2023). Consequently, we have shown that none of the existing methods for variety selection can reliably and consistently predict the agronomic performance in target environments (Figure 3, Figure 4). In contrast, NVT in Australia has been more receptive toward adopting state-of-the-art statistical advances in predicting variety performance (Smith et al 2001, 2005, 2015, 2021).

Our results suggest that the variety recommendation system can be easily improved using the GBLUP approach which can be implemented with existing trial designs. Aside from not needing any modification to the current trial designs, this approach only requires all varieties to be genotyped using a standard platform and a switch from estimating simple means to fitting mixed models. GBLUP is widely used in selection for lines and crosses in many major plant and animal breeding programs (Clark and van der Werf 2013, Hickey et al 2017). Here, we have shown that method I5B and A5B are superior to their counterparts and A5B provides the best prediction of all methods considered (Figure 2). Aside from the overall benefits, we have also demonstrated that the GBLUP approach produces slight improvement across trial sites (Figure 3), reliable and consistent results across years (Figure 4), and minimal yield deficits (Figure 6).

Improvement to the variety recommendation system can be further increased by including unreleased or candidate varieties from the National List (NL) and breeders’ lines. Additional data from more individuals and environments allows the GBLUP model to better partition genetic, environment and GEI effects (Edwards et al 2019, Voss-Fels et al 2019). Before going into the RL trials, a candidate variety has to be granted plant variety rights through evaluations for Distinctness, Uniformity and Stability (DUS) and Value for Cultivation or Use (VCU) in NL trials. Therefore, genotyping candidate varieties may serve dual purposes in applying genomic DUS in variety registration (Yang et al 2021) and GBLUP in variety recommendation. Because data confidentiality can be an issue with unreleased varieties or breeders’ lines, a recently developed homomorphic encryption method provides a straightforward way to maintain privacy in data sharing (Mott et al 2020). Furthermore, this encryption method does not require further validation as it has been shown to work well with GBLUP (Zhao et al 2024).

In addition to our GBLUP approach, there are also other methods that may improve the variety recommendation system. For example, treating variety as a random instead of fixed effect with appropriately fitted GEI terms has been shown to improve variety performance prediction (Smith et al 2005, Piepho et al 2008, Molenaar et al 2018). This method can also be described as switching of the statistical model from Best Linear Unbiased Estimation (BLUE) to Best Linear Unbiased Prediction (BLUP) for variety effect. Genomic marker data is not always necessary for this method, but it does require either fitting data for all replicates within each trial site in the one-stage analysis or carrying over residual variance-covariance matrix in the two-stage analysis (Piepho et al 2008). Unfortunately, neither of this information is available from the AHDB. BLUP aside, there is a method known as Additive Main Effects and Multiplicative Interaction (AMMI) which accounts for GEI by fitting only fixed effects in its model (Gauch 1988, Annicchiarico 1997). Similar to our observations on the challenges in using fixed over random effects, AMMI has been shown to be less accurate than BLUP (Piepho 1994). Factor Analytic Linear Mixed Model (FA-LMM) is an improved method based on AMMI where it attempts to identify similar environments and reduce model complexity (Piepho 1997, Smith et al 2015, 2021). Lastly, given recent developments in Machine Learning (ML) and Artificial Intelligence (AI), there has been many ML/AI-based methods described for applications in variety recommendation system (Newman and Furbank 2021, Balakrishnan et al 2023, Hasan et al 2023, Han et al 2024, Shams et al 2024).

RL has come a long way since its inception in 1944 and is now celebrating its 80^th^ anniversary in 2024. Our results suggest that it is timely to revamp RL by revising the statistical methods for variety recommendation. This change will benefit breeders and growers, contribute to sustainable agriculture, and tackle various threats from climate change.

## Acknowledgement

We are grateful to the IMPROMALT consortium for sharing the spring barley genomic marker data. We appreciate valuable discussion with the past and present members of the Principal’s Research Group, including Rajiv Sharma, Ian Dawson, and David Marshall. This work is supported by funding from the Scottish Society for Crop Research (SSCR) awarded to IM.

