## Supplementary materials for "Opportunities to improve the recommendation of plant varieties under the Recommended List (RL) system"

Table S1. Spring barley trial sites from 2002 to 2019.

Table S2. Spring barley varieties from 2002 to 2019.

Figure S1. Distribution of counties with spring barley trials.

Figure S2. Methods and criteria for predicting variety performance.

Figure S3. Correlations between predicted and observed yield ranks in all 29 analyzed trial sites.

### Table S1. Spring barley trial sites from 2002 to 2019.

Geographical information, numbers of varieties trialed each year and indicators of whether the site is analyzed are provided for each of the 80 spring barley trial sites from 2002 to 2019.

| Site | County | Region | Lat (°) | Long (°) | Nation | 2002 | 2003 | 2004 | 2005 | 2006 | 2007 | 2008 | 2009 | 2010 | 2011 | 2012 | 2013 | 2014 | 2015 | 2016 | 2017 | 2018 | 2019 | Analyzed |
|---|---:|---:|---:|---:|---:|---:|---:|---:|---:|---:|---:|---:|---:|---:|---:|---:|---:|---:|---:|---:|---:|---:|---:|---:|
| Tain | Highland | North | 57.812 | -4.055 | Scotland | 0 | 19 | 0 | 0 | 0 | 21 | 22 | 21 | 22 | 22 | 29 | 34 | 38 | 24 | 20 | 21 | 25 | 21 | Yes |
| Rosskeen | Highland | North | 57.696 | -4.211 | Scotland | 0 | 0 | 25 | 27 | 0 | 0 | 0 | 0 | 0 | 0 | 0 | 0 | 0 | 0 | 0 | 0 | 0 | 0 | No |
| Sandend | Aberdeenshire | North | 57.685 | -2.748 | Scotland | 0 | 0 | 0 | 0 | 0 | 0 | 0 | 0 | 0 | 0 | 0 | 0 | 38 | 24 | 20 | 0 | 25 | 21 | Yes |
| Elgin | Moray | North | 57.644 | -3.313 | Scotland | 19 | 0 | 0 | 0 | 0 | 0 | 0 | 0 | 0 | 0 | 0 | 0 | 0 | 0 | 0 | 21 | 0 | 0 | No |
| Inverurie | Aberdeenshire | North | 57.284 | -2.374 | Scotland | 19 | 19 | 25 | 27 | 0 | 21 | 22 | 21 | 0 | 0 | 0 | 0 | 0 | 0 | 0 | 0 | 0 | 0 | Yes |
| Dyce | City of Aberdeen | North | 57.203 | -2.188 | Scotland | 0 | 0 | 0 | 0 | 29 | 0 | 0 | 0 | 22 | 22 | 0 | 0 | 0 | 0 | 0 | 0 | 0 | 0 | No |
| Laurencekirk | Aberdeenshire | North | 56.832 | -2.468 | Scotland | 19 | 19 | 25 | 27 | 29 | 22 | 22 | 21 | 22 | 22 | 29 | 34 | 38 | 24 | 20 | 21 | 0 | 21 | Yes |
| Lintrathen | Angus | North | 56.679 | -3.181 | Scotland | 0 | 0 | 0 | 0 | 0 | 0 | 0 | 0 | 0 | 0 | 0 | 34 | 0 | 0 | 0 | 0 | 0 | 0 | No |
| Arbroath | Angus | North | 56.559 | -2.591 | Scotland | 0 | 0 | 0 | 0 | 0 | 0 | 0 | 0 | 0 | 0 | 0 | 0 | 0 | 0 | 0 | 21 | 25 | 21 | No |
| Coupar Angus | Perth and Kinross | North | 56.547 | -3.264 | Scotland | 19 | 19 | 25 | 27 | 29 | 21 | 0 | 0 | 0 | 0 | 0 | 0 | 0 | 0 | 0 | 0 | 0 | 0 | Yes |
| Dundee | City of Dundee | North | 56.465 | -2.971 | Scotland | 0 | 0 | 0 | 0 | 0 | 0 | 0 | 0 | 0 | 0 | 0 | 34 | 38 | 24 | 20 | 0 | 0 | 0 | Yes |
| Invergowrie | Perth and Kinross | North | 56.460 | -3.062 | Scotland | 0 | 0 | 0 | 0 | 0 | 0 | 0 | 21 | 22 | 22 | 0 | 0 | 0 | 0 | 0 | 0 | 0 | 21 | No |
| Perth | Perth and Kinross | North | 56.394 | -3.432 | Scotland | 19 | 19 | 25 | 27 | 29 | 0 | 22 | 21 | 22 | 22 | 0 | 34 | 38 | 24 | 0 | 0 | 0 | 0 | Yes |
| St Andrews | Fife | North | 56.340 | -2.791 | Scotland | 19 | 19 | 0 | 0 | 0 | 0 | 0 | 0 | 0 | 0 | 0 | 0 | 0 | 0 | 0 | 0 | 0 | 0 | No |
| Aberuthven | Perth and Kinross | North | 56.322 | -3.653 | Scotland | 0 | 0 | 0 | 0 | 0 | 0 | 0 | 0 | 0 | 0 | 0 | 0 | 0 | 0 | 0 | 0 | 25 | 0 | No |
| Kinross | Perth and Kinross | North | 56.208 | -3.423 | Scotland | 0 | 0 | 0 | 0 | 0 | 0 | 0 | 0 | 0 | 0 | 0 | 0 | 0 | 0 | 20 | 21 | 25 | 21 | Yes |
| Coaltown of Balgonie | Fife | North | 56.185 | -3.126 | Scotland | 0 | 0 | 25 | 27 | 29 | 21 | 22 | 21 | 0 | 22 | 29 | 34 | 38 | 24 | 20 | 21 | 25 | 21 | Yes |
| Kinghorn | Fife | North | 56.071 | -3.174 | Scotland | 0 | 0 | 0 | 0 | 0 | 0 | 0 | 0 | 22 | 0 | 0 | 0 | 0 | 0 | 0 | 0 | 0 | 0 | No |
| North Berwick | East Lothian | North | 56.055 | -2.719 | Scotland | 0 | 19 | 25 | 0 | 0 | 0 | 0 | 0 | 0 | 0 | 0 | 0 | 0 | 0 | 0 | 0 | 0 | 0 | No |
| Longniddry | East Lothian | North | 55.976 | -2.891 | Scotland | 0 | 0 | 0 | 27 | 29 | 0 | 0 | 0 | 0 | 0 | 0 | 0 | 0 | 0 | 0 | 0 | 0 | 0 | No |
| Stenton | East Lothian | North | 55.962 | -2.603 | Scotland | 0 | 0 | 0 | 0 | 0 | 22 | 22 | 0 | 0 | 0 | 0 | 0 | 0 | 0 | 0 | 0 | 0 | 0 | No |
| Edinburgh | City of Edinburgh | North | 55.951 | -3.195 | Scotland | 19 | 0 | 0 | 0 | 0 | 0 | 0 | 0 | 0 | 0 | 0 | 0 | 0 | 0 | 0 | 0 | 0 | 0 | No |
| East Saltoun | East Lothian | North | 55.901 | -2.840 | Scotland | 0 | 0 | 0 | 0 | 0 | 0 | 0 | 21 | 22 | 22 | 29 | 34 | 38 | 24 | 20 | 21 | 25 | 21 | Yes |
| Duns | Scottish Borders | North | 55.778 | -2.344 | Scotland | 0 | 0 | 0 | 0 | 0 | 0 | 0 | 0 | 22 | 22 | 0 | 0 | 0 | 0 | 0 | 0 | 0 | 0 | No |
| Lanark | South Lanarkshire | West | 55.674 | -3.776 | Scotland | 0 | 0 | 0 | 0 | 0 | 21 | 22 | 21 | 22 | 22 | 29 | 0 | 0 | 0 | 0 | 21 | 25 | 21 | Yes |
| Coldstream | Scottish Borders | North | 55.652 | -2.251 | Scotland | 19 | 0 | 0 | 0 | 0 | 0 | 0 | 0 | 0 | 0 | 0 | 0 | 0 | 0 | 0 | 0 | 0 | 0 | No |
| St Boswells | Scottish Borders | North | 55.570 | -2.646 | Scotland | 0 | 19 | 25 | 27 | 29 | 21 | 22 | 21 | 22 | 22 | 29 | 0 | 0 | 24 | 20 | 21 | 25 | 21 | Yes |
| Ayr | South Ayrshire | West | 55.458 | -4.629 | Scotland | 19 | 19 | 25 | 27 | 29 | 21 | 0 | 0 | 0 | 0 | 0 | 0 | 0 | 0 | 0 | 0 | 0 | 0 | Yes |
| Lockerbie | Dumfries & Galloway | West | 55.122 | -3.353 | Scotland | 0 | 19 | 25 | 27 | 0 | 0 | 0 | 0 | 0 | 0 | 0 | 0 | 0 | 0 | 0 | 0 | 0 | 0 | No |
| Annan | Dumfries & Galloway | West | 54.984 | -3.262 | Scotland | 19 | 0 | 0 | 0 | 0 | 0 | 0 | 0 | 0 | 0 | 0 | 0 | 0 | 0 | 0 | 0 | 0 | 0 | No |
| Coleraine | Londonderry | West | 55.132 | -6.666 | N Ireland | 0 | 0 | 0 | 0 | 0 | 21 | 22 | 0 | 0 | 0 | 0 | 0 | 38 | 0 | 0 | 0 | 0 | 0 | No |
| Limavady | Londonderry | West | 55.045 | -6.934 | N Ireland | 19 | 0 | 0 | 0 | 0 | 0 | 0 | 0 | 0 | 0 | 0 | 0 | 0 | 0 | 0 | 0 | 0 | 0 | No |
| Strabane | Tyrone | West | 54.827 | -7.463 | N Ireland | 0 | 19 | 25 | 27 | 29 | 22 | 0 | 21 | 0 | 22 | 0 | 34 | 38 | 24 | 20 | 0 | 0 | 0 | Yes |
| Crossnacreevy | Down | West | 54.556 | -5.849 | N Ireland | 19 | 19 | 25 | 27 | 29 | 21 | 0 | 21 | 22 | 0 | 29 | 0 | 38 | 24 | 20 | 21 | 25 | 0 | Yes |
| Ballywalter | Down | West | 54.543 | -5.484 | N Ireland | 0 | 0 | 0 | 0 | 0 | 21 | 0 | 0 | 0 | 22 | 29 | 34 | 38 | 0 | 0 | 0 | 0 | 0 | Yes |
| Abergavenny | Monmouthshire | West | 51.825 | -3.019 | Wales | 0 | 19 | 0 | 27 | 29 | 19 | 22 | 0 | 0 | 0 | 0 | 0 | 0 | 0 | 0 | 0 | 0 | 0 | Yes |
| Usk | Monmouthshire | West | 51.704 | -2.903 | Wales | 0 | 0 | 25 | 0 | 0 | 0 | 0 | 0 | 0 | 0 | 0 | 0 | 0 | 0 | 0 | 0 | 0 | 0 | No |
| Bowsden | Northumberland | North | 55.669 | -2.014 | England | 0 | 0 | 25 | 0 | 29 | 22 | 22 | 21 | 0 | 0 | 0 | 0 | 0 | 0 | 0 | 0 | 0 | 0 | Yes |
| Morpeth | Northumberland | North | 55.168 | -1.686 | England | 19 | 0 | 0 | 0 | 0 | 0 | 0 | 0 | 0 | 0 | 0 | 0 | 0 | 0 | 0 | 0 | 0 | 0 | No |
| Wintringham | North Yorkshire | East | 54.147 | -0.651 | England | 0 | 0 | 0 | 27 | 0 | 0 | 0 | 0 | 0 | 0 | 0 | 0 | 0 | 0 | 0 | 0 | 0 | 21 | No |
| Rudston | East Yorkshire | East | 54.089 | -0.324 | England | 19 | 0 | 0 | 0 | 0 | 0 | 0 | 0 | 0 | 0 | 0 | 0 | 0 | 0 | 0 | 0 | 0 | 0 | No |
| Towthorpe | East Yorkshire | East | 54.021 | -1.045 | England | 0 | 0 | 0 | 0 | 0 | 0 | 0 | 0 | 0 | 0 | 0 | 0 | 0 | 0 | 0 | 0 | 25 | 0 | No |
| Bainton | East Yorkshire | East | 53.957 | -0.533 | England | 0 | 0 | 0 | 0 | 0 | 0 | 22 | 21 | 22 | 22 | 29 | 34 | 38 | 24 | 20 | 0 | 0 | 0 | Yes |
| Sancton | East Yorkshire | East | 53.842 | -0.633 | England | 0 | 0 | 0 | 0 | 29 | 20 | 0 | 0 | 0 | 0 | 0 | 0 | 0 | 0 | 0 | 0 | 0 | 0 | No |
| Cawood | North Yorkshire | East | 53.833 | -1.131 | England | 0 | 0 | 0 | 0 | 0 | 0 | 0 | 0 | 0 | 0 | 0 | 0 | 0 | 24 | 20 | 21 | 0 | 0 | No |
| Ormskirk | Lancashire | West | 53.569 | -2.884 | England | 0 | 19 | 0 | 0 | 0 | 0 | 0 | 0 | 0 | 0 | 0 | 0 | 0 | 0 | 0 | 0 | 0 | 0 | No |
| Ince Blundell | Merseyside | West | 53.524 | -3.026 | England | 0 | 0 | 25 | 27 | 29 | 0 | 0 | 0 | 0 | 0 | 0 | 0 | 0 | 0 | 0 | 0 | 0 | 0 | No |
| Market Rasen | Lincolnshire | East | 53.385 | -0.335 | England | 0 | 0 | 0 | 0 | 0 | 0 | 0 | 0 | 0 | 22 | 0 | 0 | 0 | 0 | 0 | 0 | 0 | 0 | No |
| High Legh | Cheshire | West | 53.354 | -2.451 | England | 0 | 0 | 0 | 0 | 0 | 0 | 22 | 0 | 0 | 0 | 0 | 34 | 38 | 24 | 20 | 21 | 25 | 21 | Yes |
| Runcorn | Cheshire | West | 53.342 | -2.730 | England | 19 | 0 | 0 | 0 | 0 | 0 | 0 | 0 | 0 | 0 | 0 | 0 | 0 | 0 | 0 | 0 | 0 | 0 | No |
| Horncastle | Lincolnshire | East | 53.208 | -0.113 | England | 19 | 19 | 25 | 27 | 29 | 19 | 22 | 21 | 0 | 0 | 0 | 34 | 38 | 24 | 20 | 21 | 25 | 21 | Yes |
| Fakenham | Norfolk | East | 52.831 | 0.850 | England | 0 | 0 | 0 | 0 | 0 | 0 | 0 | 0 | 0 | 0 | 0 | 0 | 0 | 0 | 0 | 0 | 0 | 21 | No |
| Woodseaves | Staffordshire | West | 52.826 | -2.299 | England | 0 | 0 | 0 | 0 | 0 | 0 | 0 | 21 | 0 | 0 | 0 | 0 | 0 | 0 | 0 | 0 | 0 | 0 | No |
| Edgmond | Shropshire | West | 52.775 | -2.412 | England | 0 | 0 | 0 | 0 | 0 | 0 | 0 | 0 | 0 | 22 | 0 | 34 | 38 | 24 | 20 | 21 | 25 | 21 | Yes |
| Kings Lynn | Norfolk | East | 52.752 | 0.402 | England | 0 | 19 | 25 | 27 | 0 | 19 | 22 | 21 | 22 | 0 | 29 | 34 | 38 | 24 | 20 | 21 | 0 | 0 | Yes |
| Wymondham | Norfolk | East | 52.569 | 1.115 | England | 19 | 19 | 25 | 27 | 29 | 19 | 22 | 21 | 0 | 22 | 29 | 34 | 38 | 24 | 20 | 21 | 25 | 21 | Yes |
| Eye | Suffolk | East | 52.320 | 1.146 | England | 0 | 19 | 25 | 27 | 29 | 0 | 0 | 0 | 0 | 0 | 0 | 0 | 0 | 0 | 0 | 0 | 0 | 0 | No |
| Little Staughton | Bedfordshire | East | 52.250 | -0.382 | England | 0 | 0 | 0 | 27 | 0 | 0 | 0 | 0 | 0 | 0 | 0 | 0 | 0 | 0 | 0 | 0 | 0 | 0 | No |
| Rougham | Suffolk | East | 52.221 | 0.804 | England | 0 | 0 | 0 | 0 | 0 | 0 | 0 | 0 | 0 | 0 | 0 | 0 | 0 | 0 | 0 | 0 | 25 | 0 | No |
| Fulbourn | Cambridgeshire | East | 52.183 | 0.222 | England | 0 | 0 | 0 | 0 | 0 | 0 | 0 | 0 | 0 | 0 | 0 | 0 | 0 | 0 | 20 | 21 | 25 | 21 | Yes |
| Otley | Suffolk | East | 52.151 | 1.221 | England | 19 | 0 | 0 | 0 | 0 | 0 | 0 | 0 | 0 | 0 | 0 | 0 | 0 | 0 | 0 | 0 | 0 | 0 | No |
| Alkerton | Oxfordshire | West | 52.083 | -1.454 | England | 0 | 0 | 0 | 0 | 0 | 0 | 0 | 0 | 0 | 0 | 0 | 34 | 0 | 0 | 0 | 0 | 0 | 0 | No |
| Callow | Herefordshire | West | 52.005 | -2.739 | England | 0 | 0 | 0 | 0 | 0 | 0 | 0 | 0 | 0 | 0 | 29 | 34 | 0 | 24 | 20 | 21 | 25 | 21 | Yes |
| Luton | Bedfordshire | East | 51.879 | -0.420 | England | 0 | 19 | 0 | 0 | 0 | 0 | 0 | 0 | 0 | 0 | 0 | 0 | 0 | 0 | 0 | 0 | 0 | 0 | No |
| Cirencester | Gloucestershire | West | 51.718 | -1.969 | England | 0 | 19 | 25 | 27 | 0 | 20 | 22 | 21 | 22 | 22 | 0 | 0 | 0 | 0 | 0 | 0 | 0 | 0 | Yes |
| South Stoke | Oxfordshire | West | 51.541 | -1.134 | England | 0 | 0 | 0 | 0 | 0 | 0 | 0 | 0 | 22 | 0 | 0 | 0 | 0 | 0 | 0 | 0 | 0 | 0 | No |
| East Malling | Kent | East | 51.287 | 0.439 | England | 0 | 0 | 0 | 0 | 0 | 0 | 0 | 0 | 0 | 0 | 0 | 0 | 0 | 0 | 0 | 21 | 0 | 0 | No |
| Adisham | Kent | East | 51.241 | 1.189 | England | 0 | 0 | 0 | 0 | 0 | 0 | 0 | 0 | 0 | 0 | 0 | 0 | 0 | 0 | 0 | 0 | 0 | 21 | No |
| Warminster | Wiltshire | West | 51.205 | -2.181 | England | 19 | 0 | 0 | 0 | 0 | 0 | 0 | 0 | 0 | 0 | 0 | 0 | 0 | 0 | 0 | 0 | 0 | 0 | No |
| Shrewton | Wiltshire | West | 51.193 | -1.903 | England | 0 | 0 | 25 | 27 | 29 | 0 | 22 | 0 | 0 | 0 | 0 | 0 | 0 | 0 | 0 | 0 | 0 | 0 | No |
| Stockbridge | Hampshire | West | 51.117 | -1.486 | England | 0 | 0 | 0 | 27 | 0 | 20 | 22 | 21 | 22 | 0 | 29 | 34 | 38 | 24 | 20 | 21 | 25 | 21 | Yes |
| Winchester | Hampshire | West | 51.060 | -1.310 | England | 19 | 19 | 0 | 0 | 0 | 0 | 0 | 0 | 0 | 22 | 0 | 0 | 0 | 0 | 0 | 0 | 0 | 0 | No |
| Blandford Forum | Wiltshire | West | 50.857 | -2.165 | England | 0 | 19 | 0 | 0 | 0 | 0 | 0 | 0 | 0 | 0 | 0 | 0 | 0 | 0 | 0 | 0 | 0 | 0 | No |
| Exeter | Devon | West | 50.726 | -3.527 | England | 0 | 19 | 0 | 0 | 0 | 0 | 0 | 0 | 0 | 0 | 0 | 0 | 0 | 0 | 0 | 0 | 0 | 0 | No |
| Stoke Climsland | Cornwall | West | 50.547 | -4.316 | England | 0 | 0 | 0 | 0 | 0 | 0 | 0 | 0 | 0 | 0 | 0 | 0 | 0 | 24 | 20 | 0 | 0 | 0 | No |
| Wadebridge | Cornwall | West | 50.517 | -4.836 | England | 19 | 0 | 0 | 0 | 0 | 0 | 0 | 0 | 0 | 0 | 0 | 0 | 0 | 0 | 0 | 0 | 0 | 0 | No |
| Totnes | Devon | West | 50.434 | -3.686 | England | 19 | 0 | 0 | 0 | 0 | 0 | 0 | 0 | 0 | 22 | 0 | 0 | 0 | 0 | 0 | 0 | 0 | 0 | No |
| Trerulefoot | Cornwall | West | 50.406 | -4.348 | England | 0 | 0 | 25 | 27 | 0 | 0 | 0 | 0 | 0 | 0 | 0 | 0 | 0 | 0 | 0 | 0 | 0 | 0 | No |
| Plymouth | Devon | West | 50.376 | -4.143 | England | 0 | 19 | 0 | 0 | 0 | 0 | 0 | 0 | 0 | 0 | 0 | 0 | 0 | 0 | 0 | 0 | 0 | 0 | No |
| Kingsbridge | Devon | West | 50.283 | -3.777 | England | 0 | 0 | 0 | 27 | 0 | 20 | 0 | 21 | 0 | 0 | 0 | 34 | 38 | 24 | 20 | 21 | 25 | 21 | Yes |

**Table S2. Spring barley varieties from 2002 to 2019.**

All spring barley varieties (control and new) from 2002 to 2019 are listed here along with their first and last years in the Recommended List (RL) trials. Information for varieties with either 2002 as first year or 2019 as last year may not be accurate because some of these varieties may span outside of the data range.

| Variety | First | Last |
|---|---|---|
| BARKE | 2002 | 2002 |
| COLSTON | 2002 | 2002 |
| COUNTY | 2002 | 2002 |
| GLOBAL | 2002 | 2002 |
| NOVELLO | 2002 | 2002 |
| SEBASTIAN | 2002 | 2002 |
| TAVERN | 2002 | 2002 |
| VORTEX | 2002 | 2002 |
| PRESTIGE | 2002 | 2004 |
| CHALICE | 2002 | 2006 |
| PEWTER | 2002 | 2006 |
| SPIRE | 2002 | 2006 |
| CELLAR | 2002 | 2007 |
| KIRSTY | 2002 | 2007 |
| RIVIERA | 2002 | 2007 |
| STATIC | 2002 | 2007 |
| COCKTAIL | 2002 | 2009 |
| DECANTER | 2002 | 2010 |
| OPTIC | 2002 | 2013 |
| ATHENA | 2003 | 2003 |
| BERYLLIUM | 2003 | 2003 |
| DRUM | 2003 | 2003 |
| FELTWELL | 2003 | 2003 |
| TOBY | 2003 | 2003 |
| CARAFE | 2003 | 2006 |
| TROON | 2003 | 2006 |
| DOYEN | 2003 | 2008 |
| REBECCA | 2003 | 2008 |
| MACAW | 2004 | 2004 |
| MINSTREL | 2004 | 2004 |
| TOUCAN | 2004 | 2004 |
| HENLEY | 2004 | 2006 |
| POWER | 2004 | 2006 |
| TOCADA | 2004 | 2006 |
| WICKET | 2004 | 2007 |
| OXBRIDGE | 2004 | 2010 |
| NFC_TIPPLE | 2004 | 2015 |
| WAGGON | 2004 | 2015 |
| WESTMINSTER | 2004 | 2015 |
| BEATRIX | 2005 | 2005 |
| CENTURION | 2005 | 2005 |
| CRIBBAGE | 2005 | 2005 |
| HYDRA | 2005 | 2005 |
| PUTNEY | 2005 | 2005 |
| POKER | 2005 | 2006 |
| APPALOOSA | 2005 | 2008 |
| PRAGUE | 2006 | 2006 |
| TAPHOUSE | 2006 | 2006 |
| TARTAN | 2006 | 2009 |
| PUBLICAN | 2006 | 2011 |
| QUENCH | 2006 | 2014 |

| Variety | First | Last |
|---|---|---|
| KNIGHTSBRIDGE | 2007 | 2007 |
| MALTBY | 2007 | 2007 |
| SNAKEBITE | 2007 | 2007 |
| JOLIKA | 2007 | 2008 |
| SWEENEY | 2007 | 2008 |
| SCOUT | 2007 | 2009 |
| BELGRAVIA | 2007 | 2017 |
| BERLIOZ | 2008 | 2008 |
| CONCHITA | 2008 | 2008 |
| VIRGIL | 2008 | 2008 |
| CROPTON | 2008 | 2010 |
| FORENSIC | 2008 | 2011 |
| CONCERTO | 2008 | 2019 |
| BENCHMARK | 2009 | 2009 |
| CAIRN | 2009 | 2009 |
| MIRAGE | 2009 | 2009 |
| YARD | 2009 | 2009 |
| GARNER | 2009 | 2014 |
| PROPINO | 2009 | 2019 |
| CHECKMATE | 2010 | 2010 |
| CROMWELL | 2010 | 2010 |
| MARIONETTE | 2010 | 2010 |
| SY_TABERNA | 2010 | 2010 |
| PANTHER | 2010 | 2011 |
| SUMMIT | 2010 | 2012 |
| SHUFFLE | 2010 | 2013 |
| MOONSHINE | 2010 | 2014 |
| BOGART | 2011 | 2011 |
| SY_ABOYNE | 2011 | 2011 |
| SY_BARRELL | 2011 | 2011 |
| SY_UNIVERSAL | 2011 | 2011 |
| CHRONICLE | 2011 | 2013 |
| OVERTURE | 2011 | 2014 |
| ODYSSEY | 2011 | 2017 |
| ACCLAIM | 2012 | 2012 |
| MAGELLAN | 2012 | 2012 |
| MOMENTUM | 2012 | 2012 |
| PINOCCHIO | 2012 | 2012 |
| SPARKLE | 2012 | 2012 |
| CROONER | 2012 | 2014 |
| GLASSEL | 2012 | 2014 |
| KWS_ORPHELIA | 2012 | 2014 |
| MONTOYA | 2012 | 2014 |
| NATASIA | 2012 | 2014 |
| RHYNCOSTAR | 2012 | 2014 |
| KELIM | 2012 | 2015 |
| TESLA | 2012 | 2015 |
| SANETTE | 2012 | 2016 |
| ALVESTON | 2013 | 2013 |
| ARTISAN | 2013 | 2013 |
| KERSTIN | 2013 | 2013 |

| Variety | First | Last |
|---|---|---|
| MALT_JAGGER | 2013 | 2013 |
| MARY | 2013 | 2013 |
| MELIUS | 2013 | 2013 |
| RENAISSANCE | 2013 | 2013 |
| KWS_AURELIA | 2013 | 2014 |
| SHALOO | 2013 | 2014 |
| SHADA | 2013 | 2015 |
| HACKER | 2013 | 2018 |
| KWS_IRINA | 2013 | 2019 |
| DRAGOON | 2014 | 2014 |
| INVICTUS | 2014 | 2014 |
| MILFORD | 2014 | 2014 |
| PATHFINDER | 2014 | 2014 |
| PIPER | 2014 | 2014 |
| RGT_CONQUEST | 2014 | 2014 |
| DEVERON | 2014 | 2015 |
| VAULT | 2014 | 2015 |
| OCTAVIA | 2014 | 2017 |
| SCHOLAR | 2014 | 2018 |
| OLYMPUS | 2014 | 2019 |
| RGT_PLANET | 2014 | 2019 |
| SIENNA | 2014 | 2019 |
| ORIGIN | 2015 | 2016 |
| OVATION | 2015 | 2018 |
| KWS_SASSY | 2015 | 2019 |
| LAUREATE | 2015 | 2019 |
| ACORN | 2016 | 2016 |
| DIOPTRIC | 2016 | 2017 |
| LG_OPERA | 2016 | 2017 |
| CHANSON | 2016 | 2019 |
| FAIRING | 2016 | 2019 |
| LG_TOMAHAWK | 2017 | 2018 |
| LG_DIABLO | 2017 | 2019 |
| RGT_ASTEROID | 2017 | 2019 |
| ACCURANCE | 2018 | 2018 |
| EMBRACE | 2018 | 2018 |
| LG_GODDESS | 2018 | 2018 |
| RGT_ORBITER | 2018 | 2018 |
| SY_CONTOUR | 2018 | 2018 |
| SY_DOLOMITE | 2018 | 2018 |
| SY_KAILASH | 2018 | 2018 |
| SY_STANZA | 2018 | 2018 |
| COSMOPOLITAN | 2018 | 2019 |
| BARBARELLA | 2019 | 2019 |
| FIREFOXX | 2019 | 2019 |
| LG_FURLONG | 2019 | 2019 |
| LG_SERENGETI | 2019 | 2019 |
| PROSPECT | 2019 | 2019 |
| RGT_SLIPSTREAM | 2019 | 2019 |
| SY_SPLENDOR | 2019 | 2019 |
| SY_TUNGSTEN | 2019 | 2019 |

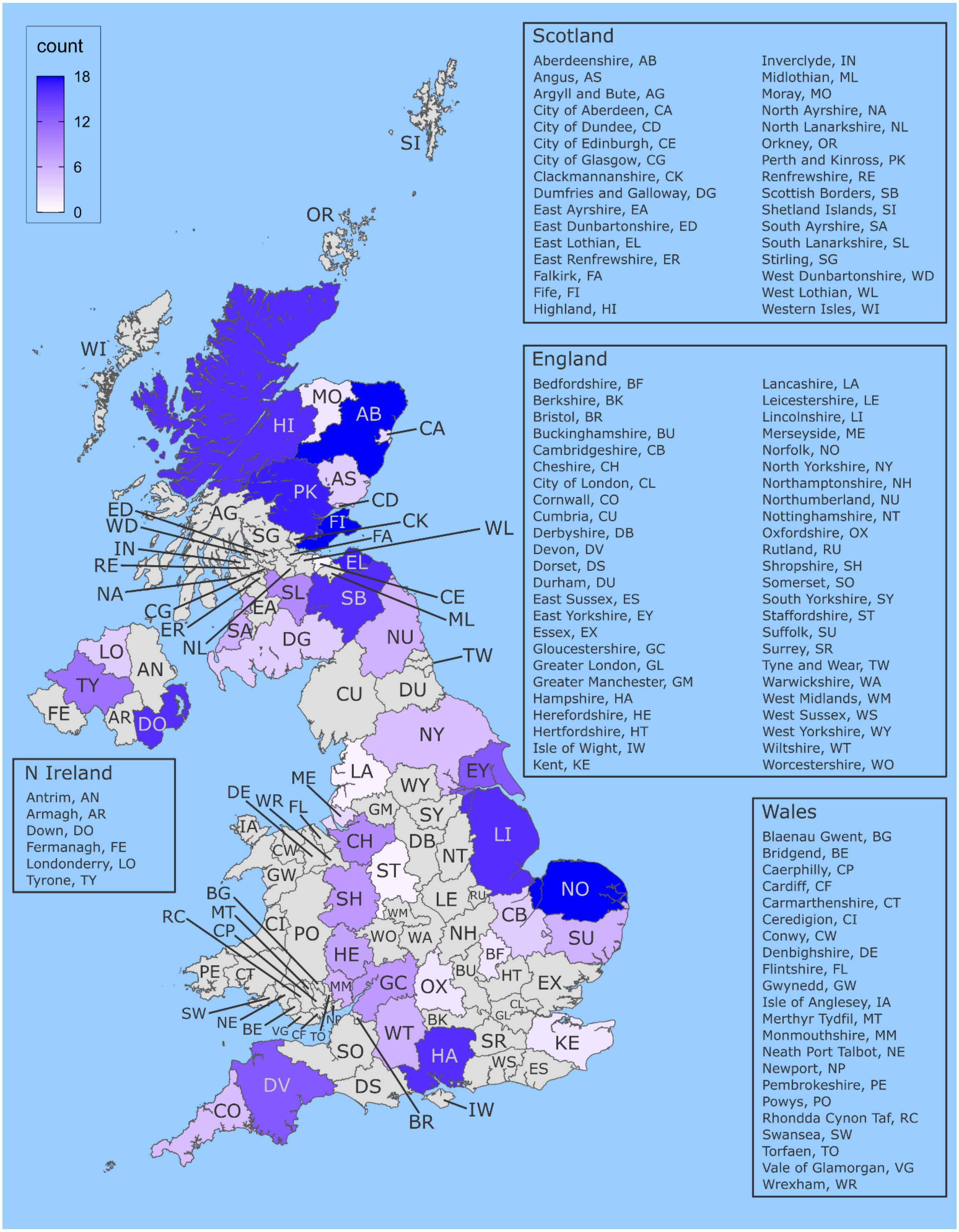


**Figure S1. Distribution of counties with spring barley trials.**

For each county, the numbers of years from 2002 to 2019 with spring barley trials are counted. Multiple trial sites per county-year contributes a maximum of one count per county-year.

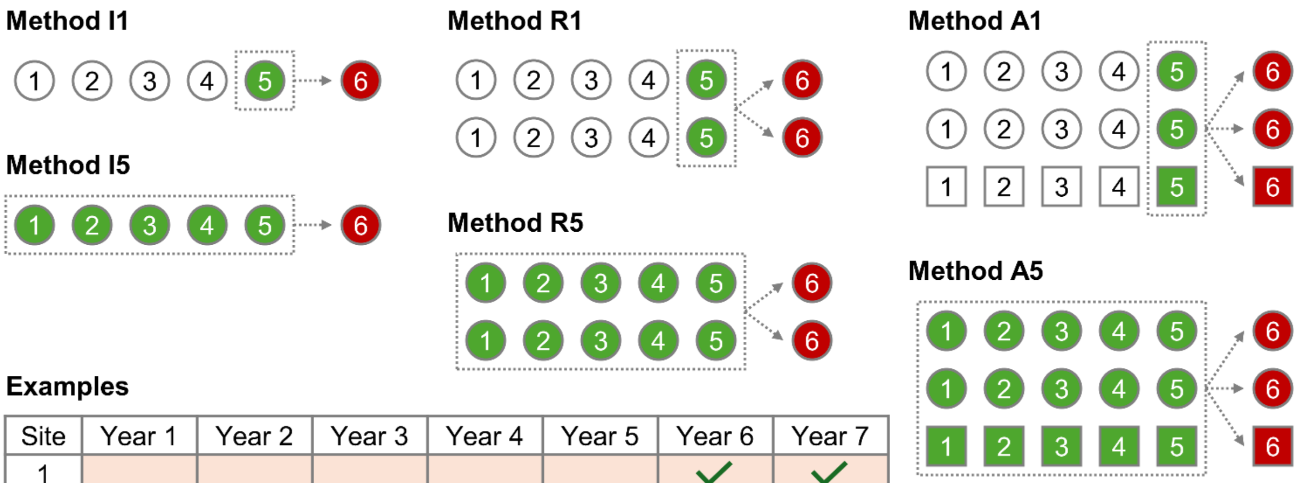


**Examples**

| Site | Year 1 | Year 2 | Year 3 | Year 4 | Year 5 | Year 6 | Year 7 |
|---|---|---|---|---|---|---|---|
| 1 | | | | | | ✓ | ✓ |
| 2 | | | | | | ✓ | ✗ 1 |
| 3 | | | | | | ✗ 2 | ✓ |
| 4 | | | | | | ✗ 3 | ✓ |
| 5 | | | | | | ✗ 1 | ✗ 2 |
| 6 | | | | | | ✗ 3 | ✗ 1 |
| 7 | | | | | | ✗ 1,2,3 | ✗ 1,2,3 |
| 8 | | | | | | ✗ 2,3 | ✗ 1,3 |
| 9 | | | | | | ✓ | ✓ |

**Data criteria for analysis**

1. Target year
2. Previous 1 year
3. 3 out of 5 previous years

**Figure S2. Methods and criteria for predicting variety performance.**

In the current AHDB variety selection tool, there are six different methods for calculating yield means for predicting future performance. Broadly, these methods use either one or five years of yield data, and calculate individual site (I), regional (R) or overall (A) means. Each row indicates a site, and each shape represents a region (AHDB classifies regions as North, West and East, see Figure 1B). Even though the dataset starts from 2002, the first target year that we can predict is 2007 as method I5, R5 and A5 use five years of data. Because variety trials are often sparse and the same site is not always used every year, three additional criteria were set to maximize usable trial data. Examples of how these criteria work are provided here with fully met criteria indicated by green ticks and failed criteria annotated along with red crosses.

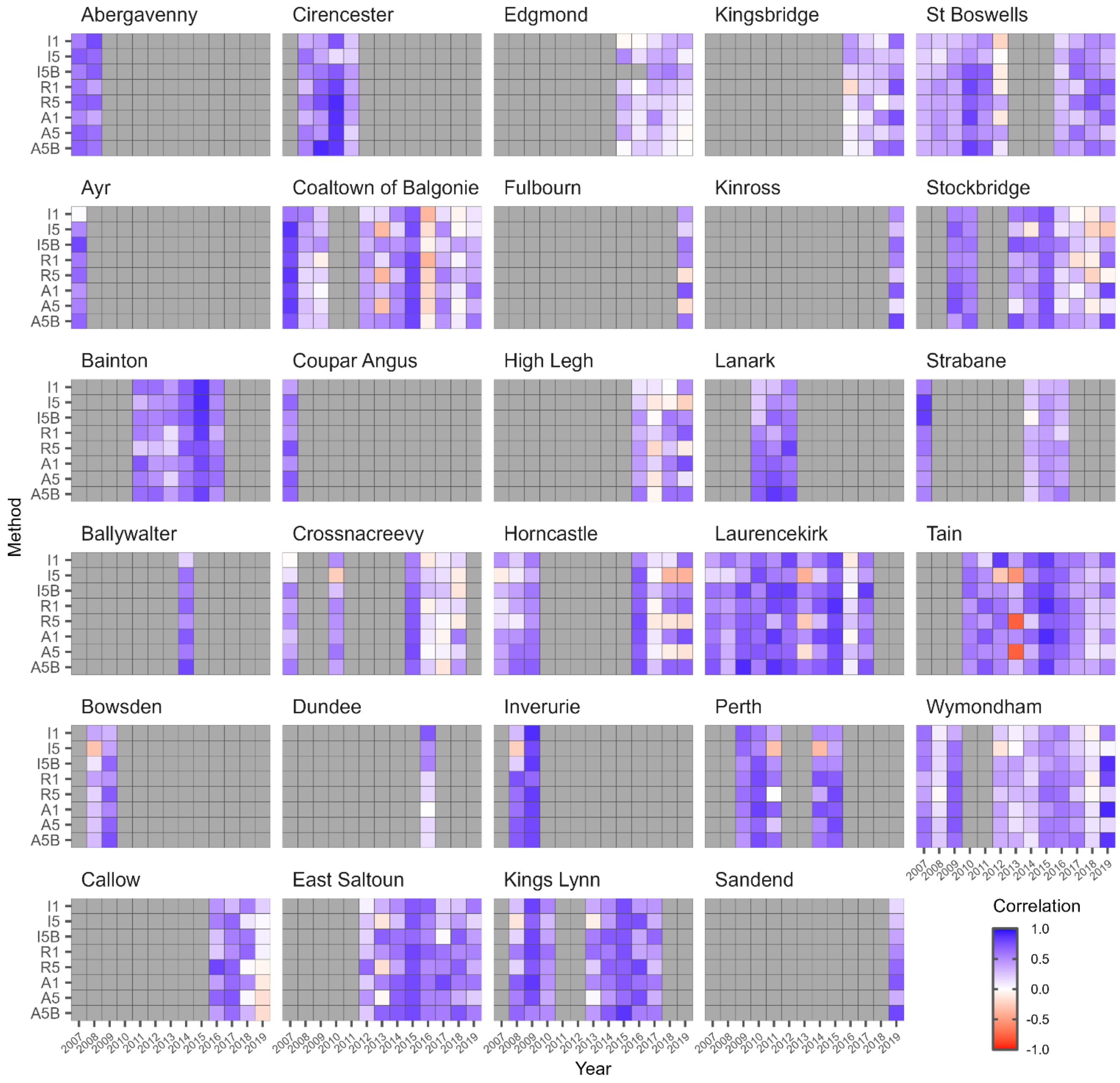


**Figure S3. Correlations between predicted and observed yield ranks in all 29 analyzed trial sites.**

Heatmap of correlations for eight methods and 13 years. A condensed version of this figure with six trial sites with the most data (Coaltown of Balgonie, Laurencekirk, St Boswells, Stockbridge, Tain, Wymondham) can be found in Figure 3A.
